# Detection of Japanese Encephalitis Virus in Mosquitoes Collected from Selected Areas of Laguna, Philippines Using an Optimized One-Step RT-PCR Protocol

**DOI:** 10.64898/2026.09.03.749309

**Authors:** John Leonard R. Chan, Bernadette C. Mendoza, Lucille C. Villegas, Joseph S. Masangkay, Loinda R. Baldrias, Thaddeus M. Carvajal, Kozo Watanabe

## Abstract

Japanese encephalitis virus is a leading cause of encephalitis in humans and agricultural animals across Southern and Eastern Asia. It is primarily transmitted by mosquitoes of the *Culex* genus, notably *Culex tritaeniorhynchus*. In the Philippines, JEV is reported to account for 7% to 18% of clinical cases of meningitis and encephalitis. Therefore, updated and comprehensive data on circulating JEV strains in the country are crucial for developing effective mitigation strategies. This study aimed to optimize an assay for JEV detection using one-step reverse transcription polymerase chain reaction (RT-PCR). Additionally, the presence of JEV in mosquitoes collected from eleven cities and municipalities in Laguna was assessed, and the mosquito carriers were identified based on morphological characteristics. Results demonstrated that the optimized one-step RT-PCR assay successfully detected JEV at various dilutions, with a limit of detection (LoD) of 0.115 ng/μL. The assay exhibited 100% specificity, with no amplification observed for any viruses other than JEV. Sequence analysis of the positive control, JEV SA14-14-2 RNA, revealed 96.96% to 98.41% similarity to JEV reference sequences in the NCBI database. Among the 5,522 female mosquitoes collected, *Culex tritaeniorhynchus* and *Culex gelidus* were the most prevalent species. The highest mosquito species diversity and the most even distribution were found in the three barangays of Paete, but Siniloan, Laguna had the highest vector density for the three *Culex* species, *Culex tritaeniorhynchus*, *Culex gelidus*, and *Culex fuscocephala*. RT-PCR analysis of 189 pooled mosquito samples revealed no detectable JEV genetic material. This study is likely the first investigation of JEV presence in mosquito samples from Laguna, Philippines. These findings contribute to proactive JEV surveillance, enhancing our understanding of the virus’s presence and distribution. The results underscore the importance of targeted prevention and control measures to mitigate JEV transmission in high-risk areas.

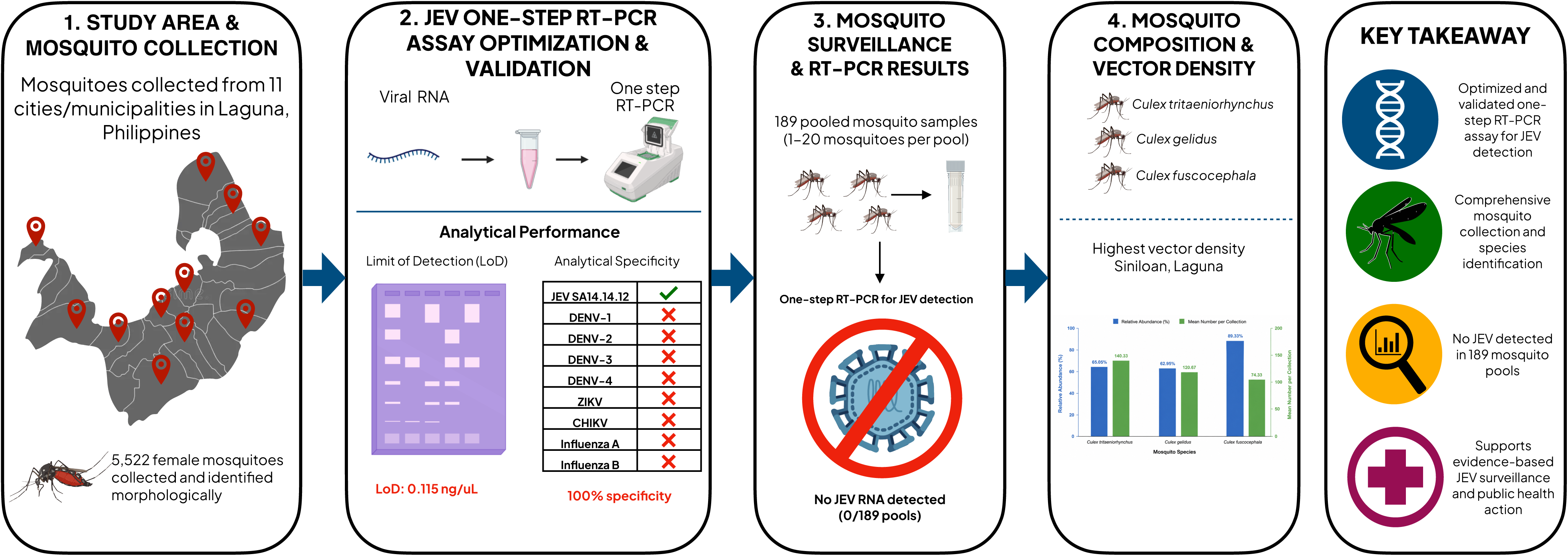

## INTRODUCTION

Japanese Encephalitis Virus (JEV), the causative agent of Japanese encephalitis (JE), belongs to the family *Flaviviridae*, genus Flavivirus. JEV is related to some other clinically important Flaviviruses, like dengue virus, tick-borne encephalitis virus, and yellow fever virus.^1^ It was first reported in Japan during the 19th century and was isolated in monkeys during the early 1930s by passaging in mice. The recent viruses causing encephalitis in humans in several parts of the world like North America were shown to be related to but distinct from JEV.^2^

Japanese encephalitis is considered endemic in Asia, and various cases have been reported in the Philippines, China, India, Pakistan, South Korea, Japan, and in other parts of Southeast Asia. It is the major cause of encephalitis in both humans and agricultural animals in southern and eastern Asia.^3^ The World Health Organization (WHO) includes JEV as one of the important emerging pathogens, approximately 3 billion people are at risk of Japanese encephalitis infection, particularly in areas with reported cases. Every year, around 20,000 confirmed cases occur with 6,000 fatalities. The fatality rate ranges between 5% to 30%, but up to 50% of survivors of the disease may experience neurological problems, such as intellectual disability, while approximately one-third achieve full recovery.

In the Philippines, the Japanese encephalitis virus is the leading cause of acute encephalitis in humans. In 2019, approximately 15% of cases of acute encephalitis were attributed to JEV.^4^ There were eighteen (18) clinical studies from 1972 to 2013 that documented 257 cases of laboratory-confirmed JE with 7-18% reported as combined clinical meningitis and encephalitis, and 16-40% as clinical encephalitis cases.^5^ The majority of individuals affected by JE are children under the age of 15, with a mortality arte of 6-7%.^5^ In August 2017, it was reported that there were 133 confirmed cases of Japanese encephalitis in the country, resulting in nine deaths.^6^

Five genotypes of Japanese encephalitis virus (JEV) have been identified based on nucleotide sequences from the capsid/pre-membrane and envelope regions.^7^ Predominantly, genotypes I and III occur in epidemic regions, whereas genotypes II and IV are associated with endemic disease. It has been postulated that differences in strain virulence may explain the clinical epidemiology.^8^ The most recently published study on JEV in the Philippines, conducted in San Jose, Tarlac, from May 2009 to July 2010, revealed that out of 28,700 mosquitoes tested, JEV genotype III was detected in *Culex tritaeniorhynchus*.^9^ However, more up-to-date information is needed to improve our understanding of the current epidemiological situation and to develop effective mitigation strategies in the Philippines.

This study aimed to optimize a one-step RT-PCR assay for JEV detection and to determine its possible presence in mosquitoes collected from selected areas in Laguna, Philippines. The findings may contribute to JEV surveillance using molecular assays and provide insights into the virus’s distribution within mosquito populations. These data can inform policy development and guide the implementation of effective prevention and control strategies for JE infection and other mosquito-borne diseases. Furthermore, findings from this study, along with future research, may serve as a basis for potentially integrating JE vaccination into the national immunization program.

## MATERIALS AND METHODS

Study Area, Sample Determination, and Sample Collection The study area covered selected parts of Laguna, Philippines, which were chosen due to the presence of key elements in the JEV transmission cycle: swine farms, rice fields, and residential areas. Samples were randomly collected from various Laguna locations, including Los Baños, Pagsanjan, Paete, Sta. Cruz, Nagcarlan, Calauan, Siniloan, San Pedro, Santa Rosa, San Pablo, and Cabuyao (Figure *1*). Mosquito traps were provided by De La Salle University Biological Control Research Unit, Biñan City, Laguna, and their locations were recorded using Google Maps. Sampling occurred between June and November 2019, aligning with the typical peak months of JE cases in the Philippines, which are July to October.^10^

**Figure 1:**
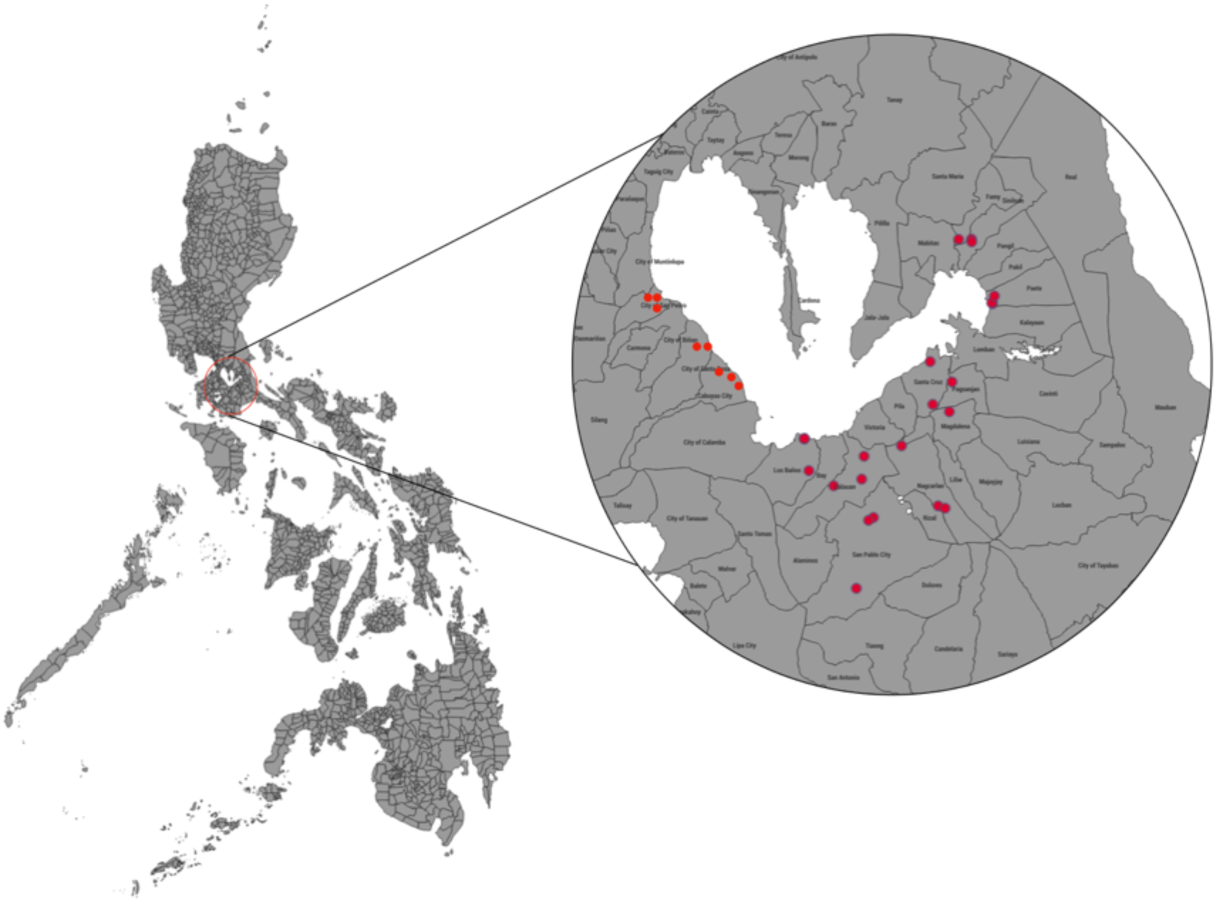
The map of the province of Laguna, Philippines showing the sites of collection of the mosquito samples. Selected municipalities and cities were each assigned to have three (3) barangays to be collected upon within June to November 2019.

The CDC Miniature Light Trap Model 512 (John W. Hock Company, Gainesville, FL) was utilized for mosquito collection. These traps were positioned at a short distance from swine farms, suspended at a height of 5-6 feet above the ground, and away from artificial light sources. Collection took place from 17:00H to 6:00H the following day.

Only female mosquitoes were collected and grouped based on tentative species identification, with a maximum of 20 mosquitoes per tube. A representative sample from each group was separated and placed in microcentrifuge tubes for species confirmation. Species confirmation was performed using a dichotomous key method^11–12^ and involved consultation with entomologists from the Medical Entomology Laboratory at the Research Institute for Tropical Medicine. All pooled samples were preserved in RNA*later*^TM^ (Thermo Fisher Scientific, USA) to maintain nucleic acid integrity. Vector abundance was calculated by tallying the total number of mosquitoes of a specific species collected and dividing it by the number of traps used per night.^13^ For species diversity indices, the Simpson’s Index of Diversity (1-D), Shannon Diversity Index (H’), and Evenness Index (E) were used to assess mosquito species diversity in each barangay.

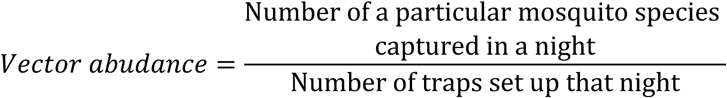

### Viral Controls

The JEV SA14-14-2 live-attenuated virus, used in the vaccine manufactured by the Chengdu Institute of Biological Products, served as the positive control and was obtained from the Virology Laboratory, Research Institute for Tropical Medicine (RITM).

Other viral nucleic acids, including Dengue virus (serotypes I–IV), Chikungunya virus, Zika virus, H1N1, AH3, and Influenza B strains, were obtained from the Advanced Molecular Technologies Laboratory, RITM, for assay specificity determination.

Detection of JEV in the Collected Samples Using One-step RT-PCR *Nucleic acid extraction.* Mosquito pooled samples, along with the positive controls for JEV were subjected to nucleic acid extraction using the MagMax™ Viral/Pathogen Nucleic Acid Isolation Kit (Thermo Fisher Scientific, USA) on the KingFisher™ Flex Purification System. After removing RNA*later^TM^* (Thermo Fisher Scientific, USA), 400 µL of sample was combined with 530 µL of Binding Solution, followed by the addition of 20 µL of magnetic beads, as per the manufacturer’s instructions. The extraction process involved four plates: Wash 1 Plate (1,000 µL wash buffer), Wash 2 Plate (1,000 µL 80% ethanol), Wash 3 Plate (500 µL 80% ethanol), and Elution Plate (80 µL elution solution). Additionally, 10 µL of proteinase K was added to the sample plate, which contained the Binding Solution, samples, and magnetic beads. The KingFisher™ Flex Purification System was programmed as MVP_Flex. Finally, 80 µL of eluate was collected, and 5 µL was transferred to microcentrifuge tubes for quantification using a DS-11 Series Spectrophotometer/Fluorometer (DeNovix, USA) *Optimization of the One-step RT-PCR Assay for Detection of JEV Envelope Gene.* A one-step RT-PCR assay was employed to detect the envelope gene using a JEV-specific primer set (Table 1).^14–15^ Modifications were made to the PCR master mix to optimize detection. The protocol was adopted and replicated using the JEV SA14-14-2 strain as a positive control. Among the modifications implemented were the inclusion of a complementary DNA (cDNA) synthesis step at 55°C for 30 minutes and an adjustment of the annealing temperature to 56°C for 30 seconds within the PCR profile. These modifications aimed to enhance the amplification efficiency and improve the detection of the JEV envelope gene during the one-step RT-PCR procedure.

**Table 1.**
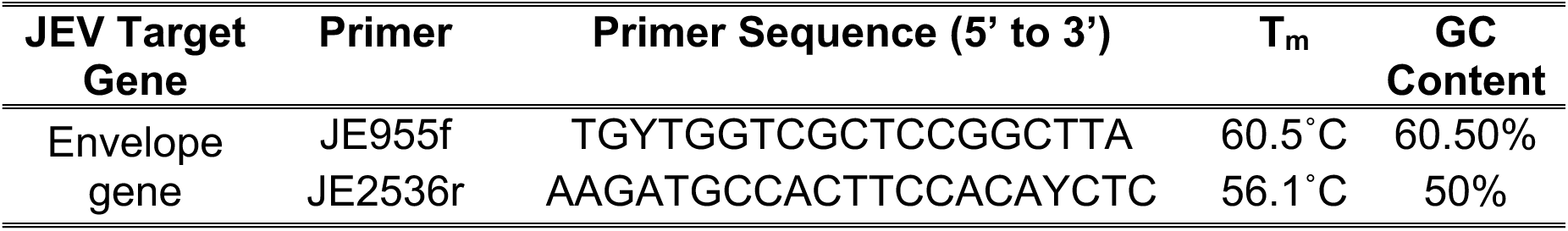
Oligonucleotide primers for the detection of Japanese Encephalitis Virus (JEV) ^15^

#### JEV Detection using the Optimized One-step RT-PCR Assay

Total nucleic acid (TNA) extracted from pooled mosquito samples served as the template for one-step RT-PCR to detect JEV. The JEV SA14-14-2 live-attenuated virus nucleic acid extracts were used as positive controls, while molecular-grade water served as the negative control. The envelope gene was amplified using a specific primer pair¹⁵ (Table 1) and the SuperScript™ III One-Step RT-PCR System with Platinum™ Taq DNA Polymerase (Invitrogen, Thermo Fisher Scientific, USA). The primers targeted a 1,561-bp product, validated for compatibility with four JEV genotypes (GenBank: LC461960.1, AF045551.2, AY316157.1, L48961.1) using SnapGene software. The PCR was performed using a modified thermal cycler program, and amplification products were analyzed via gel electrophoresis on a 1.5% agarose gel. Visualization was achieved using a 1:100 dilution of SYBR™ Safe DNA gel stain (Invitrogen, Thermo Fisher Scientific, USA).

#### Limit of Detection (LOD)/Analytical Sensitivity of the Optimized One-Step PCR Protocol

To determine the initial concentration of JEV nucleic acid in the positive control material, quantification was performed using the Qubit™ Flex Fluorometer and the Qubit™ RNA High Sensitivity (HS) Assay Kit (Thermo Fisher Scientific, USA). A serial 10-fold dilution to extinction of the JEV SA14-14-2 strain (positive control) was then prepared. To establish the lowest detectable concentration of the target analyte using the optimized JEV One-Step RT-PCR assay, one microliter from each dilution was used as the template. Each dilution was tested in triplicate using the optimized JEV One-Step RT-PCR assay.

#### Sequencing of the Envelope Gene of the Japanese Encephalitis Virus SA14-14-2 Live-Attenuated Virus

Amplicons generated from stock and serial dilutions (10⁻¹ to 10⁻¹⁰) of the positive control (JEV SA14-14-2) used in the limit of detection experiment were subjected to Sanger sequencing to verify the analytical sensitivity results. The samples were purified using the PureLink® PCR Purification Kit (Thermo Fisher Scientific, USA) to remove non-specific bands. Purified samples were analyzed via 1.5% agarose gel electrophoresis (AGE) and visualized using a 1:100 dilution of SYBR™ Safe DNA gel stain. Cycle sequencing was performed using the BigDye™ Terminator v3.1 Cycle Sequencing Kit (Applied Biosystems, Foster City, CA, USA), with JEV SA14-14-2 included as a positive control. Sequencing reaction purification was carried out using the BigDye™ XTerminator™ Purification Kit (Applied Biosystems, Foster City, CA, USA), followed by loading of the purified samples into a 3730xl Genetic Analyzer for sequencing.

#### Multiple Alignments

The nucleotide sequences of the E gene from positive samples and the positive control were compared with JEV isolates representing each genotype from the National Center for Biotechnology Information (NCBI) GenBank database. Sequence analysis was performed using Molecular Evolutionary Genetics Analysis 11 (MEGA 11) software. Multiple sequence alignments were conducted using the Clustal W program to determine the percentage similarity between the aligned sequences.

#### Analytical Specificity of the Optimized One-Step PCR Protocol

The optimized one-step RT-PCR assay for JEV was evaluated for cross-reactivity with other arboviruses that may be present in the samples, including Dengue virus serotypes I–IV, Chikungunya virus, and Zika virus, as well as three influenza viruses (H1N1, AH3, and Influenza B), which serve as representatives outside the arbovirus group. To assess analytical specificity, 1 µL of RNA extract from each virus-positive control material was used as a template.

## RESULTS

### Mosquito Samples

The study was conducted from June to November 2019, beginning in Siniloan, Laguna, Philippines. The total count of female mosquito samples collected from eleven municipalities and cities (Table 2.) Species identification was performed using a dichotomous key.¹¹⁻¹² A total of 5,522 mosquitoes were collected from seven municipalities (Los Baños, Pagsanjan, Siniloan, Paete, Nagcarlan, Sta. Cruz, and Calauan) and four cities (San Pablo, Cabuyao, San Pedro, and Sta. Rosa). The majority of mosquitoes were collected from Cabuyao, Laguna (21.41%, N=5,522), followed by Siniloan, Laguna (20.75%, N=5,522). Mosquitoes from seven areas were tentatively identified to species level through morphological analysis using specific body parts. However, specimens from Calauan, San Pedro City, Cabuyao, and Sta. Rosa City (53.80%, N=5,522) could not be identified to species due to missing morphological features. Despite this limitation, these samples were included in the analysis and were labeled based on their collection sites. Nine mosquito species were identified: *Culex vishnui, C. gelidus, C. tritaeniorhynchus, C. quinquefasciatus, C. fuscocephala, Aedes aegypti, A. albopictus, A. poicilus, Mansonia uniformis,* and *Toxorhynchites* sp. The data on vector abundance of each identified species (Table 3), this measurement represents the relative number of mosquitoes during the sampling period.^13^

**Table 2.**
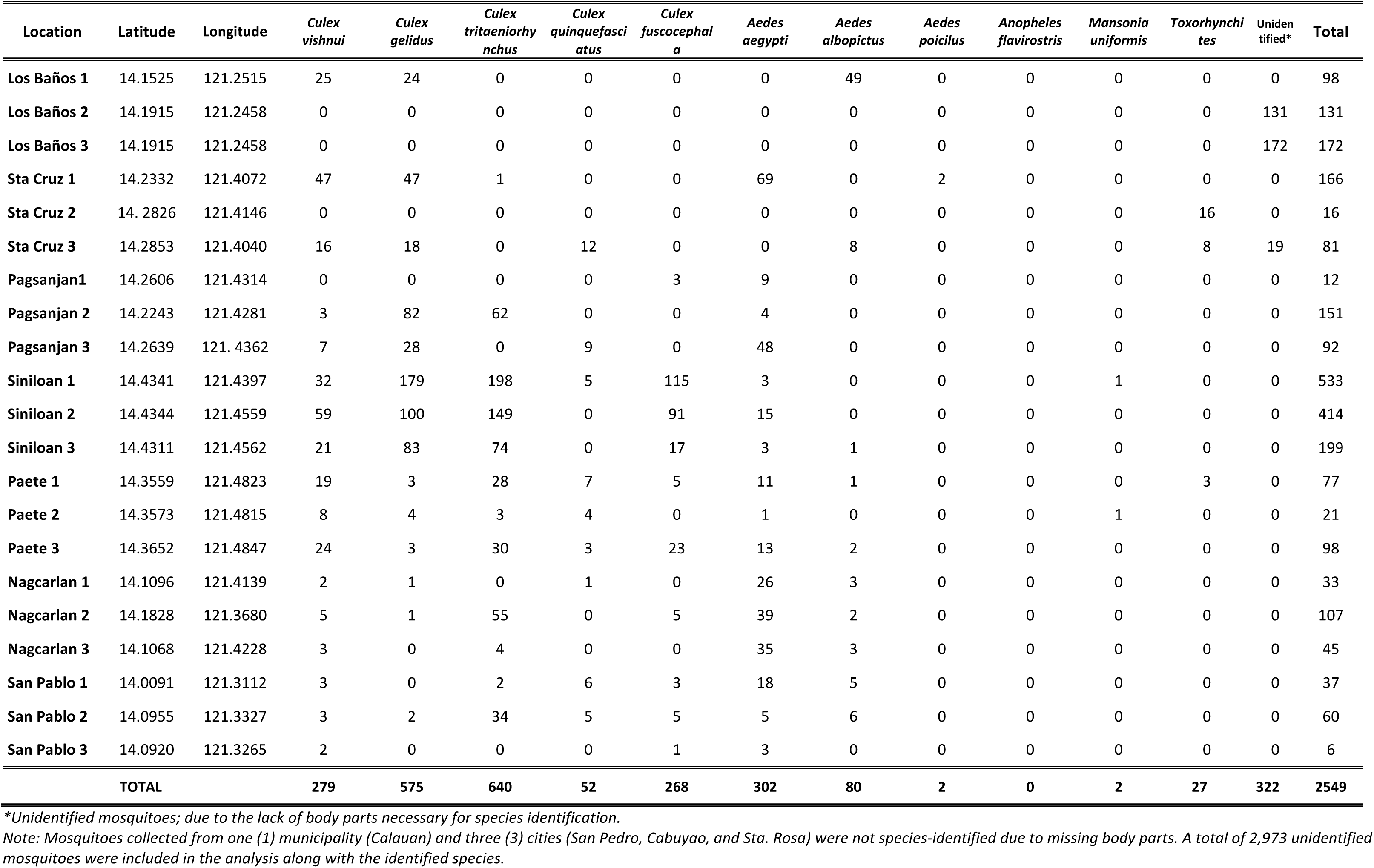
The location and latitude of the sampling sites, and the number of mosquito species per selected municipality and city in Laguna, Philippines collected from June to November 2019: putatively identified using the dichotomous key.^11–12^

**Table 3.**
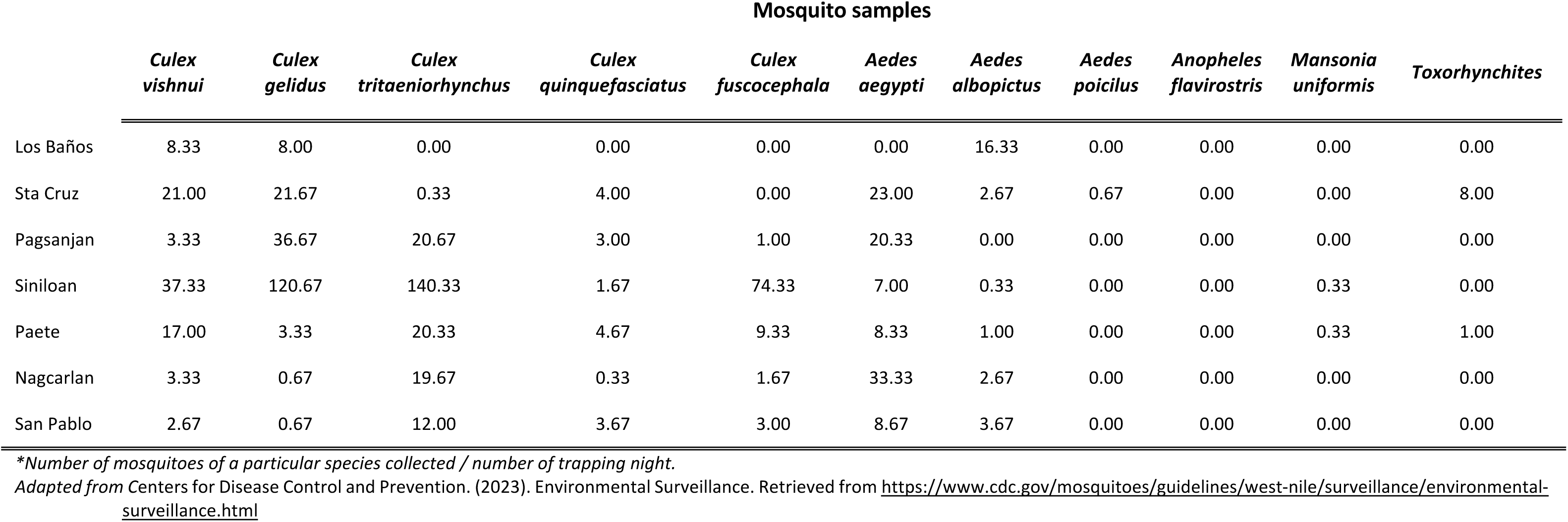
Vector abundance* per mosquito species collected in seven selected cities and municipalities in Laguna, Philippines

The majority of mosquito species with high vector density were found in Siniloan, Laguna, possibly due to the marked presence of rice fields. The predominant species identified were *Culex tritaeniorhynchus* (65.05%; 140.33), *C. gelidus* (62.95%; 120.67), *and C. fuscocephala* (89.33%; 74.33). The Shannon Diversity Index (H’) and Simpson’s Index of Diversity (1-D) indicated that Paete 1 (H’ = 1.70; 1-D = 0.70) had the highest mosquito diversity, followed by two other barangays in Paete, Laguna: Paete 3 (H’ = 1.61; 1-D = 0.70) and Paete 2 (H’ = 1.57; 1-D = 0.73). In contrast, Los Baños 2 and 3 (H’ = 0.00; 1-D = 0.50) and Sta. Cruz 2 (H’ = 0.00; 1-D = 0.52) exhibited no mosquito diversity, suggesting the dominance of a single species. Additionally, Paete 2 (E = 0.81) had the most even distribution of mosquito species among all barangays (Table 4). Mosquito samples from the four other cities and one municipality were excluded from the analysis due to unidentified species.

**Table 4.**
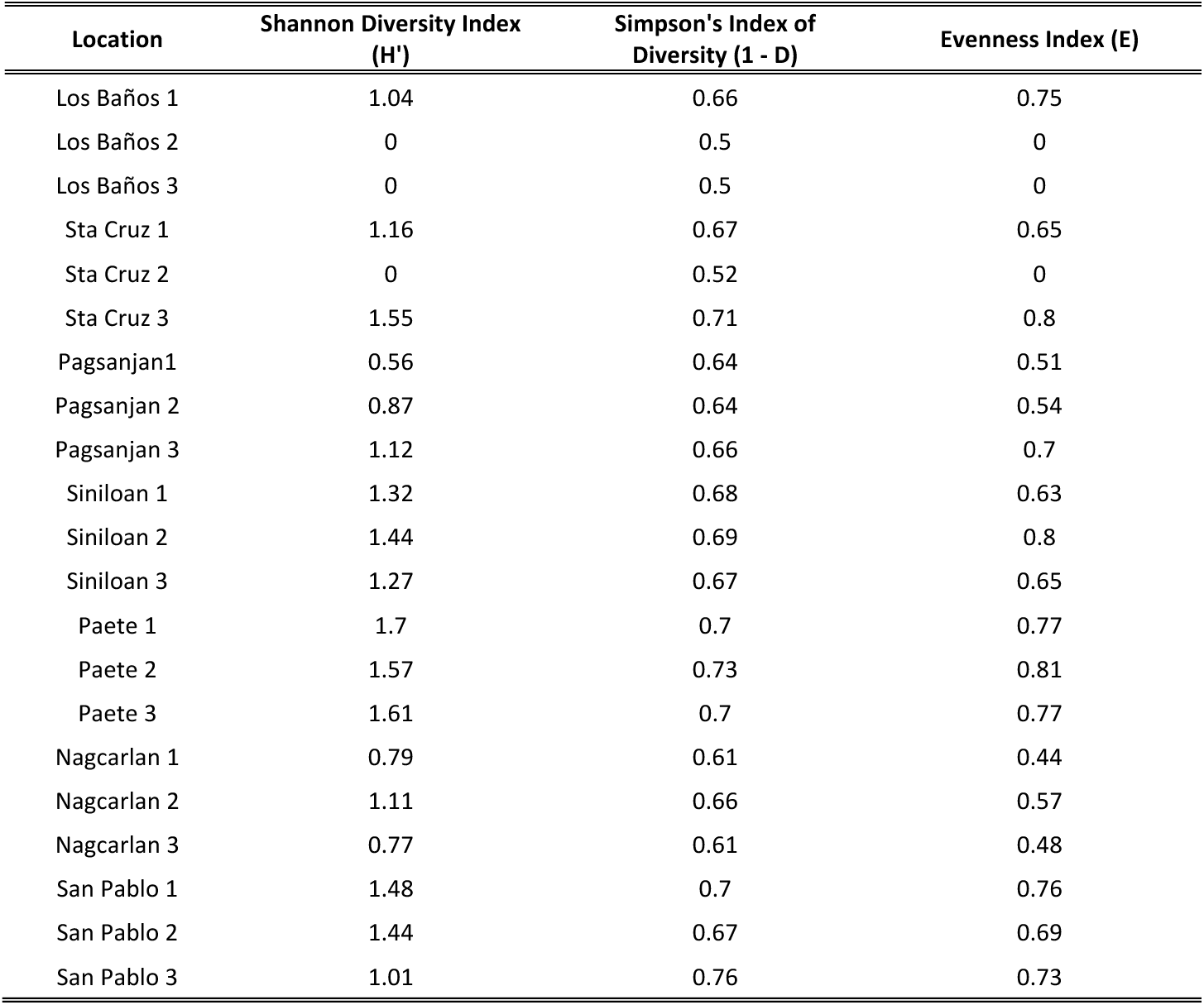
Diversity indices of mosquitoes at seven cities and municipalities in Laguna, Philippines from June to November 2019.

Optimization and Technical Performance Validation of a One-Step Reverse Transcription PCR for Japanese Encephalitis Virus Detection A positive control was used to optimize the one-step reverse transcription PCR (RT-PCR) assay for JEV. This served as the template material for assay optimization and was obtained from the Virology Laboratory at the Research Institute for Tropical Medicine. The positive control consisted of the JEV SA14-14-12 strain, derived from the live attenuated SA14-14-2 (CD-JEV) vaccine. To enhance assay performance, the PCR profile was modified by incorporating an additional incubation step at 55°C for 30 minutes to facilitate cDNA synthesis. Additionally, the annealing temperature was adjusted. The effects of these modifications on JEV detection (Figure 2A). SnapGene software v6.0 analysis confirmed that the primer set matched all four previously sequenced JEV genotypes. This finding validated the primers’ in-silico sensitivity and their capability to detect multiple JEV genotypes.

**Figure 2.**
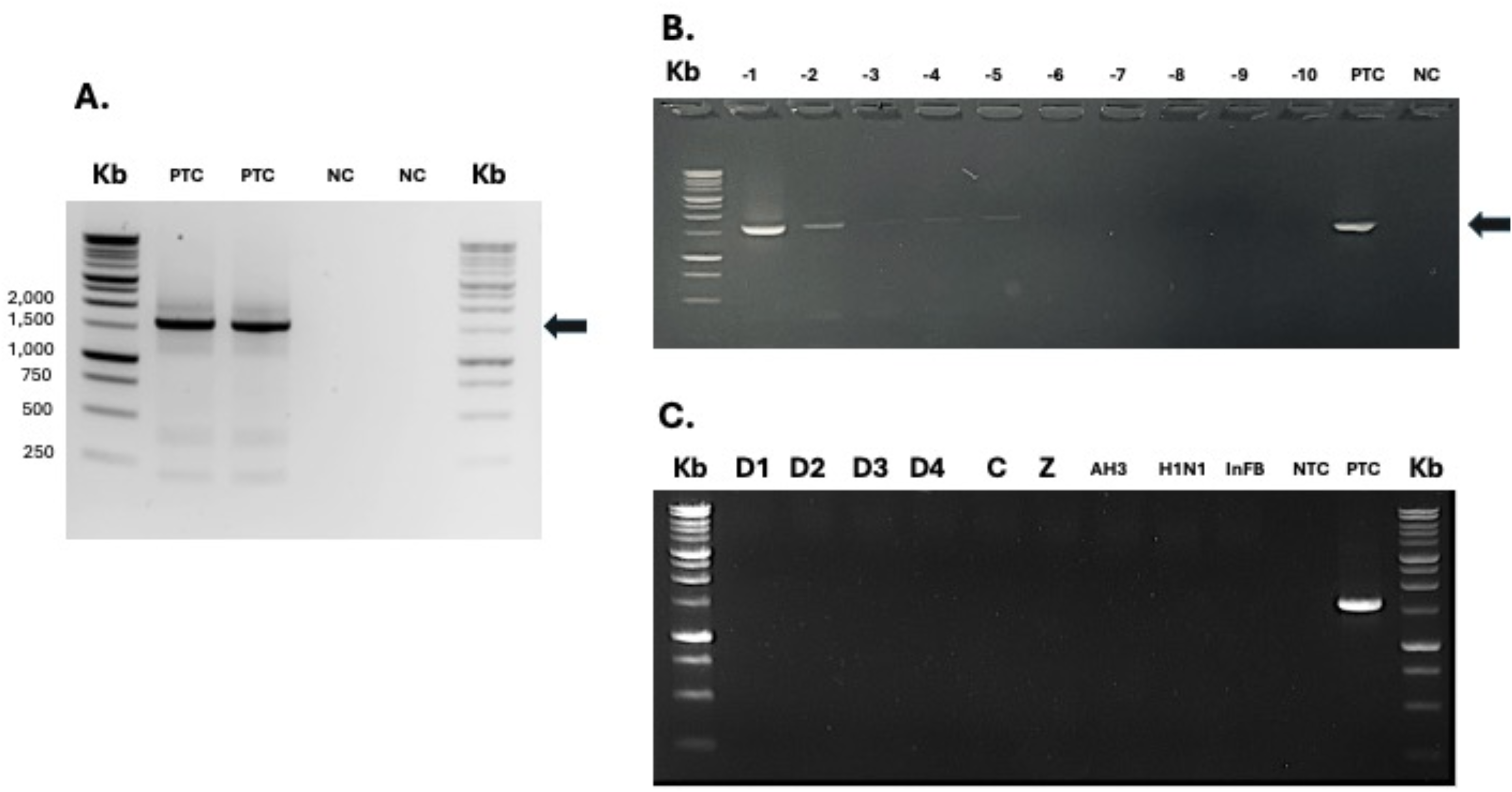
A: Electrophoretic analysis of JEV SA 14.14.12. strain as the positive control with an approximate band size of 1500bp. Kb: Molecular weight ladder; PTC: Positive Control; NC: Negative Control. **B:** Limit of detection of optimized one-step RT-PCR for the envelope (E) gene of JEV using a 10-fold dilution of the JEV SA14.14.12. strain RNA extract (positive control). PTC, Positive Control; NC: Negative Control; Lanes -1 to -10 represent 10-fold dilution. **C:** Specificity of the optimized one-step RT-PCR assay for envelope (E) gene of JEV. D1-4: four serotypes of Dengue virus; C: Chikungunya virus; Z: Zika virus; AH3 and H1N1: Influenza A; Influenza B; PTC: positive control (JEV SA14.14.12. strain); NTC: no template control/negative control.

#### Analytical Sensitivity / Limit of Detection of the Optimized One-Step PCR for Japanese Encephalitis Virus Detection

The analytical sensitivity of the optimized JEV one-step RT-PCR assay was evaluated by determining the limit of detection through a 10-fold serial dilution of the JEV positive control material. The initial RNA extract concentration was 10.40 ng/μL; however, some dilutions fell below the detection range of the Qubit™ RNA High Sensitivity (HS) assay (0.1–20 ng/μL), indicated as “STL” (sample out of range, too low). Agarose gel electrophoresis (Figure 2B) revealed visible PCR amplicons from the 1/10 (10⁻¹) to the 1/100,000 (10⁻⁵) dilution, demonstrating the assay’s sensitivity. The number of genomic copies per microliter (copies/μL) was calculated using the Bio-Synthesis formula: dividing the single-stranded RNA concentration (ng/μL) by the RNA molecular weight (g/mol) and multiplying by Avogadro’s number.^16^ The JEV positive control at 10.4 ng/μL corresponded to approximately 1.22 × 10¹⁰ copies/μL of JEV SA14- 14-2 RNA. The RNA concentrations at various dilutions were as follows: 1.42 ng/μL (10⁻¹), 0.7 ng/μL (10⁻²), 0.44 ng/μL (10⁻³), 0.129 ng/μL (10⁻⁴), and 0.115 ng/μL (10⁻⁵).

#### Analytical Specificity of the Optimized One-Step PCR for Japanese Encephalitis Virus Detection

The specificity experiment, using the designated virus panel, confirmed that the optimized assay exhibited no cross-reactivity with non-target analytes. It successfully amplified and detected only JEV SA14-14-2 nucleic acid material, demonstrating 100% analytical specificity for the tested flaviviruses and influenza viruses (Figure 2C). To further validate the assay’s specificity, future studies should incorporate additional viruses and bacteria commonly found in mosquitoes, providing a more comprehensive assessment of its specificity against a broader spectrum of potential pathogens.

#### Sequences of the Envelope Gene of Japanese Encephalitis Virus

To further validate the results of the optimized One-Step RT-PCR assay, sequence characterization of the JEV E gene amplicons was performed using Sanger sequencing. Sequence analysis confirmed the accuracy of the One-Step RT-PCR method. NCBI-BLAST analysis of the positive control template material revealed a nucleotide sequence similarity of 96.96% to 98.41% with known JEV sequences. The BLAST search results identified the closest match as the Japanese encephalitis virus isolate SA14-14-2-PHK17 (Accession number: MH258850.1), indicating a high degree of similarity in the E gene region and further corroborating the specificity of the assay.

Detection of JEV in the Collected Mosquito Samples using One-step RT-PCR Following the successful optimization of the one-step reverse transcription PCR assay for JEV detection, a total of 189 pooled mosquito samples collected from selected areas in Laguna, Philippines, were tested. The results indicated that JEV was not detected in any of the analyzed mosquito pools (Figure 3).

**Figure 3.**
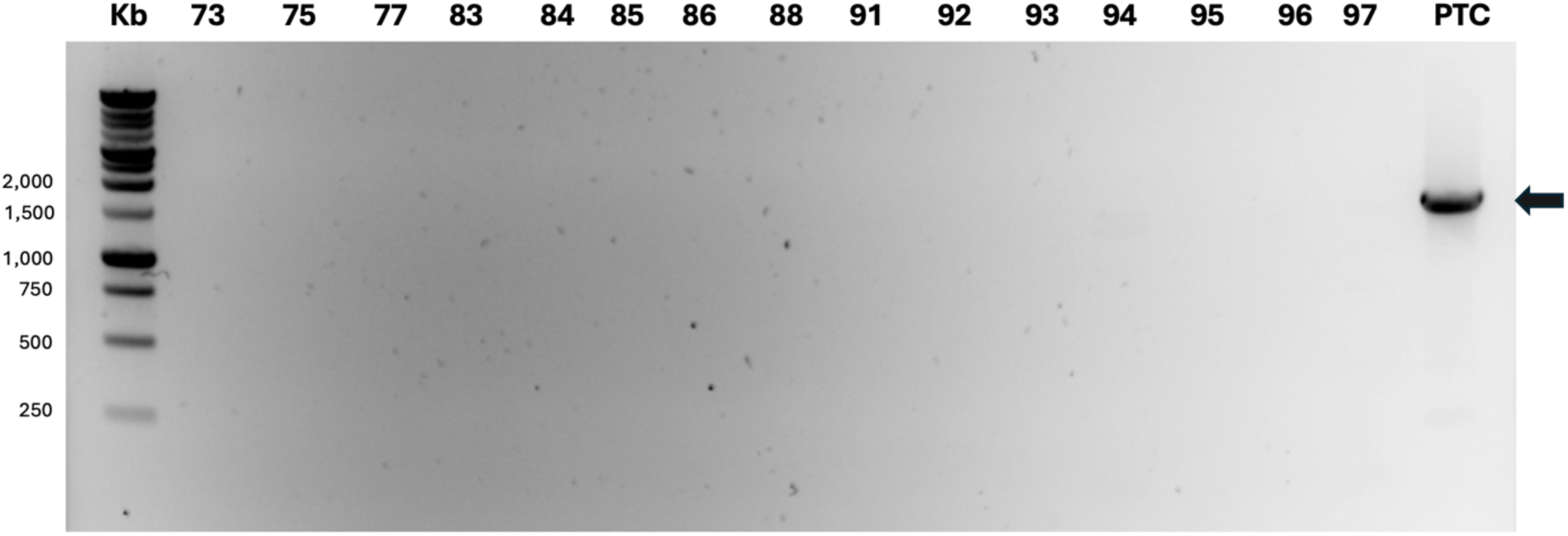
Agarose gel electrophoresis analysis of one-step reverse transcription PCR for Japanese encephalitis virus detection in some of the mosquito samples collected from Paete and Siniloan, Laguna, Philippines. Kb: DNA Molecular Ladder; Lane 2-16, mosquito samples from Paete and Siniloan, Laguna, Philippines; Lane 17, JEV SA14.14.2 (Positive Control).

## DISCUSSION

In the Philippines, mosquito surveillance efforts related to zoonotic diseases primarily focus on dengue and malaria. However, it is important to note that many mosquito species have the potential to be infected by multiple arboviruses and act as vectors, including Zika, Chikungunya, and Japanese encephalitis viruses. This study also identified vector mosquito species, such as *A. aegypti, A. albopictus, A. poicilius, M. uniformis,* and *Toxorhynchites.* Notably, *A. poicilius* was found to be a vector for filariasis. Despite the implementation of the National Filariasis Elimination Program (NFEP) by the Department of Health, 33 cases of lymphatic filariasis were reported in South Cotabato, Philippines last 2021.^17^

In general, the study found that *Culex* species were abundant, with three of them acting as vectors for the Japanese encephalitis virus (JEV). Specifically, *C. tritaeniorhynchus, Culex vishnui,* and *C. gelidus* are the main vectors for JEV, although theoretically, other genera such as *Aedes, Anopheles*, and *Mansonia* can also transmit JEV.^18^ The *Culex* species, particularly *C. tritaeniorhynchus*, is widely distributed across Southeast Asia and other tropical countries.^19^

The highest mosquito species diversity and the most even distribution of mosquito species were found in the three barangays of Paete, Laguna (average H’ = 1.63; E = 0.71), but Siniloan, Laguna had the highest vector density for the three *Culex* species. The environmental conditions of the seven cities and municipalities, including rice paddies and swine farms, provide favorable conditions for the proliferation of *Culex* species. In the JEV transmission cycle, the presence of swine as amplifying hosts, along with the prevalence of rice paddies and human activity, plays a crucial role. From 2009 to 2010, JEV genotype 3 was identified in *Culex tritaeniorhynchus* samples from Tarlac province^9^, suggesting potential JEV circulation despite the geographical distance from Laguna. High mosquito densities in risk assessment areas are often associated with arbovirus outbreaks; however, elevated mosquito populations do not always result in virus transmission, as the vector infection rate remains a critical factor.^13^ Understanding vector abundance is essential for planning and monitoring mosquito control efforts and evaluating the effectiveness of arbovirus mitigation strategies

Two *Aedes* species, *A. aegypti and A. albopictu*s, known vectors of the dengue virus, were also found in various areas of Laguna. In March 2023, the World Health Organization reported a substantial increase in dengue cases in the Philippines, highlighting the importance of continuous vector surveillance and control measures.^20^ Furthermore, the detection and isolation of the JEV in *A. albopictus* in Malaysia and Taiwan suggest its potential role as a secondary vector.^18^ Studies have demonstrated that *A. albopictus* can transmit JEV with a low infectious blood meal dose, indicating that mosquitoes in this study, including non-*Culex* species, may contribute to JEV transmission.^21^ Regular monitoring of vector abundance is essential for the effective control of potential arbovirus outbreaks in Laguna and across the country

Establishing the accuracy of the RT-PCR assay was essential for reliable JEV detection. In this study, the assay demonstrated high sensitivity, as a diluted positive control containing less than 0.115 ng/μL of JEV RNA yielded a positive result using the one-step RT-PCR. Similarly, a study tested four different JEV isolates using real-time quantitative RT-PCR (rt-qPCR) with 1 ng/μL of RNA as the template. Their assay generated a sigmoidal amplification curve with a cycle quantification (Cq) value of <40.0, indicating that RT-PCR can detect even low concentrations of viral RNA.²²

In addition, the specificity of the optimized one-step reverse transcriptase PCR (RT-PCR) assay for JEV was evaluated. The results demonstrated that no amplification was observed with other arboviruses, including all four serotypes of dengue virus (DENV), chikungunya virus (CHIKV), and Zika virus (ZIKV), as well as three respiratory viruses: H1N1, AH3, and Influenza B. Establishing the specificity of the assay was crucial to ensure that amplification and detection were exclusively targeted toward the JEV pathogen.

Sequencing of the E gene from the PCR product of the JEV positive control, using the optimized one-step RT-PCR assay, further confirmed the reliable detection of JEV. The sequence obtained exhibited a high similarity of 96% to 98% when compared to Japanese encephalitis virus sequences available in the NCBI database. This level of sequence similarity further reinforced confidence in the accurate identification of JEV by the assay.

The absence of JEV detection during the sample collection period in Laguna may suggest the effectiveness of existing mitigation efforts against arboviruses in the region. In response to the Dengvaxia scare, the Dengue Task Force was activated in 2019 to address dengue and other arbovirus transmissions in CALABARZON.^23^ Various strategies were implemented, including the four key components of the enhanced 4S strategy: Search and destroy mosquito breeding sites, Secure self-protection, Seek early consultation, and Support misting/spraying.^24^ These control measures may have contributed to reducing the presence of JEV in mosquito populations. Several other factors may also explain the negative results. One possibility is the timing of mosquito collection. In the Philippines, JE cases tend to peak annually from July to October, coinciding with the wet season. From 2014 to 2017, JE cases were reported year-round, but the highest incidence was observed during these months.^10^ Consequently, the collection period for this study was selected based on these peak months. However, this approach may have introduced sampling bias by excluding other months, such as January to May and December, when the virus may still be present but at lower levels. This seasonal variation could have contributed to the apparent non-detection of JEV in mosquitoes collected in Laguna, Philippines.

It is important to recognize that the absence of detected JEV does not necessarily indicate the complete absence of the virus in the area. Migratory birds play a crucial role in JEV transmission, particularly herons and egrets, which are known reservoirs of the virus. Monitoring the migration patterns of these birds could help identify optimal timeframes for sample collection to enhance JEV detection.^25^ Findings from this study may serve as a basis for recommending the implementation of JEV surveillance among migratory bird populations in the Philippines.

The findings of this study highlight the need for enhanced efforts in determining and monitoring the circulating genotype of JEV in the country. Identifying the specific genotype in circulation is crucial for selecting and administering the appropriate JE vaccine to high-risk populations. Given that different JEV genotypes may exhibit variations in antigenic properties, genotype-specific information is essential for developing targeted vaccination strategies and improving the effectiveness of JE control programs.

Furthermore, the locally optimized RT-PCR assay for JEV detection presented in this study addresses the existing gap in molecular diagnostic tools for JEV identification from mosquito samples. This assay contributes to the broader goal of mitigating JE in the Philippines by supporting the implementation of evidence-based measures for arbovirus prevention and control.

## CONCLUSION

Japanese encephalitis virus is a major cause of acute encephalitis worldwide and the leading cause in the Philippines. While *Culex* species are the primary vectors of JEV, several studies have demonstrated that *Aedes* species are also susceptible to the virus. In this study, various mosquito species were collected, with *Culex tritaeniorhynchus* being the most prevalent. The highest mosquito vector density was observed in Siniloan, Laguna. The optimized one-step reverse transcriptase PCR assay demonstrated high sensitivity and specificity for JEV detection. Using this assay, 189 mosquito pools collected from Laguna between June and November 2019 were examined, and no JEV was detected. These findings suggest the absence of circulating JEV in Laguna during the study period. Possible explanations include effective arbovirus mitigation efforts by local health authorities. Continued surveillance of circulating JEV genotypes is essential to monitor potential reintroduction or emergence of the virus and to inform vaccination strategies in Laguna, Philippines. This study provides valuable insights into the absence of JEV during the specific collection period and underscores the importance of ongoing surveillance and preventive measures against JEV.

## ACKNOWLEDGMENTS

The main author would like to express his gratitude to the CHED-Philippine- California Advance Research Institute and DOST-SEI for their support during his master’s degree studies. To the colleagues at the Research Institute for Tropical Medicine are acknowledged, namely Lei Lanna M. Dancel, Timothy John R. Dizon, Amalea Dulcene Nicolasora, Othoniel John Onza, Stephen Paul Ortia, Joanna Ina Manalo, Francisco Polotan, Miguel Abulencia, Aileen May Mojica, Inez Andrea Medado, Criselda Bautista, Ava Kristy Sy-Lee, and Mary Ann Ammugauan for their invaluable technical support during the conduct of the research.

